# Transporter-Mediated Uptake of Microcystin-LR in Human Trophoblasts: Regulation By Oxygen Concentration and Cell Fusion

**DOI:** 10.64898/2026.03.22.713491

**Authors:** Michael Jeffrey Campbell, Mokshitkumar Patel, Chenghui Jiang, Xia Wen, Shuo Xiao, Lauren M. Aleksunes

**Affiliations:** Rutgers University, Joint Graduate Program in Toxicology, 170 Frelinghuysen Road, Piscataway, NJ 08854, USA; Rutgers University, Ernest Mario School of Pharmacy, Department of Pharmacology and Toxicology, 170 Frelinghuysen Road, Piscataway, NJ 08854, USA; Environmental and Occupational Health Sciences Institute, 170 Frelinghuysen Rd, Piscataway, NJ 08854, USA

**Keywords:** microcystin, OATP, placenta, hypoxia, cell fusion, protein phosphatase

## Abstract

**Background:** Rising global temperatures and eutrophication are increasing the intensity and frequency of cyanobacterial harmful algal blooms that release toxins including microcystin-LR (MC-LR). MC-LR inhibits protein phosphatases in the human liver and brain, but its accumulation in the placenta is unclear. Placental transporter expression varies across pregnancy and is influenced by physiological cues, such as low oxygen concentrations which activate HIF1A, and trophoblast cell fusion forming syncytiotrophoblasts that engage CREB-driven transcription. This study examined whether MC-LR accumulates in placental cells, which transporters mediate uptake, and how these transporters are regulated by HIF1A and CREB.

**Methods:** Intracellular accumulation of MC-LR (0.1–10 µM, 3 hour) was measured in human cytotrophoblasts (JAR, BeWo) and extravillous trophoblasts (HTR-8/SVneo) by western blotting for MC-LR–adducted proteins. Organic anion transporting polypeptide (OATP) involvement was tested using cyclosporin A (10 µM), an OATP inhibitor, before exposure to the OATP substrate or MC-LR. Cells were also cultured under 3%, 8%, or 20% O₂ to induce hypoxic responses or treated with forskolin (a potent intracellular cAMP inducer) to stimulate cell fusion before MC-LR exposure.

**Results:** MC-LR accumulated in all three placenta cell lines in a concentration-dependent manner. Cyclosporin A reduced MC-LR uptake by 57% in JAR cells, confirming OATP-mediated transport. Low O_2_ increased OATP4A1 expression and function but reduced protein phosphatase expression, decreasing MC-LR–bound proteins by 52–72%. Forskolin increased OATP4A1 expression and enhanced MC-LR uptake >2.5-fold.

**Conclusion:** MC-LR enters placental trophoblasts via active OATP transport, likely OATP4A1, and uptake increases under hypoxia and trophoblast fusion.

## Introduction

Rising temperatures and related anthropogenic actions, including the use of nitrogen-rich fertilizer, have accelerated eutrophication and increased the frequency and intensity of harmful algal blooms (HABs), warranting the closure of lakes and ponds.^1–3^ As the incidence of HABs continues to expand, research regarding human exposure to HAB toxins is greatly important for risk characterization.^4; 5^ Microcystins (MCs), the most common and widely distributed HAB toxins, are a class of monocyclic peptide toxins produced by several cyanobacterial genera and have been detected across freshwater and marine-influenced environments.^6; 7^ Humans are exposed to MCs through a variety of routes, including oral, inhalation, and dermal.^8^ The provisional guideline values for drinking water set by the World Health Organization limit MC concentrations to 0.3 μg/L for children and 1 μg/L for adults.^9^ In a study of 2,804 lakes in the United States, 18% and 9.9% of lakes had concentrations above the drinking water guideline for children and adults, respectively.^2^ Recent data in humans confirm that internal exposure to MCs is occurring. For example, 720 subjects in a cross-sectional study in Hunan Province, China, had detectable concentrations of MCs in their blood (0.02-0.6 ug/L, median concentration: 0.15 ug/L).^10^ Currently, there is limited biomonitoring of MCs, which has hampered our understanding of how widespread exposure may be. During HAB bloom events, environmental concentrations of MCs can increase dramatically, with toxin exposure levels reported to be hundreds to thousands fold higher than drinking-water or recreational health advisory values.^11–13^

MCs primarily accumulate in the liver, kidney, and reproductive organs through organic anion transporting polypeptides (OATPs). As members of the solute carrier (SLCO) superfamily, OATP transporters are located in the plasma membrane and control the uptake, and in some instances the efflux, of nutrients and xenobiotics.^14^; ^15^ OATPs are sodium-independent, pH-dependent, active uptake transporters that are multispecific for a variety of substrates, including bile acids, hormones, and drugs.^16; 17^ These proteins are ubiquitously expressed across organ systems, with unique isoforms enriched in different tissues. Prior studies have reported tissue- and isoform-specific OATP-mediated uptake of MCs. For example, OATP1B1 and OATP1B3 are responsible for the uptake of MCs into human hepatocytes.^18–20^ Additional OATP isoforms have been identified as capable of MC transport, including OATP1A2 in the blood-brain barrier, revealing the need to investigate other OATP isoforms for their ability to transport MCs in other tissues.^21; 22^ Once inside cells, MCs covalently bind and inhibit serine/threonine protein phosphatase 1 (PP1) and 2A (PP2A) as well as generate reactive oxygen species, damage DNA, and stimulate apoptosis.^23^; ^24^ Additional protein targets covalently modified by MCs, beyond PPs, have also been identified. These include direct inhibition of proteasome subunits D4/B9 and disruption of cellular metabolism through inhibition of pyruvate kinase 2M.^25–27^ These additional protein targets may contribute to the cellular toxicity of MC-LR.

Growing evidence indicates that the placenta is a biologically relevant and vulnerable target of MCIZLR, as demonstrated by both *in vivo* and *in vitro* models. Studies in mice have demonstrated that prenatal exposure to MC-LR (5 or 20 μg/kg) from gestational day 13 to 17 disrupts the structure of the placental vasculature and reduces fetal weight.^28; 29^ Moreover, *in vitro* exposure of human extravillous trophoblast cells to MC-LR at concentrations of 1-3 μM for 24 hours impaired trophoblast migration and invasion, two hallmark responses associated with preeclampsia.^28; 30^ These data support the ability of MC-LR to enter trophoblasts. Bulk RNAseq data from the Pregnancy Outcome Prediction Study Placentaome Database of healthy, term placenta villous tissue revealed rich expression of *SLCO2A1, SLCO2B1*, and *SLCO4A1* transcripts.^31^ Likewise, western blotting has demonstrated enrichment of OATP1A2, 2A1, 3A1, and 4A1 on the apical surface and OATP2B1, 4A1, and 5A1 on the basolateral membranes of human placental tissue.^32^ These mounting data suggest that the placenta is a target of MC toxicity, although the exact ability of trophoblast cells to transport and accumulate MC-LR is unknown.

Gestational changes in oxygen tension engage transcriptional signaling pathways that alter placenta transporter expression, creating developmental windows in which MC uptake may be enhanced. In early pregnancy, the maternal spiral arteries are occluded by extravillous trophoblasts, creating a naturally low oxygen environment of about 3% O_2_ which activates hypoxia-inducible factors (HIFs) that regulate genes involved in angiogenesis, metabolism, and transporter expression.^37^ As pregnancy progresses, arterial remodeling increases maternal blood flow, raising oxygen concentration to ∼8-10% in later trimesters.^33–36^ Researchers often model the oxygen environment of placental development using controlled atmosphere chambers, with 3% O₂ representing early gestation and 8% O₂ reflecting mid-to-late trimesters.^38^ Human BeWo cytotrophoblasts (CTBs) cultured under conditions of low oxygen exhibit significant upregulation of glucose transporters (GLUT) as a result of HIF1A signaling.^39^ These changes in placenta transporter expression highlight that oxygen concentration may be a key determinant of susceptibility to MC toxicity across pregnancy.

Throughout pregnancy, cytotrophoblasts proliferate and fuse into multinucleated syncytiotrophoblasts (STB) at the maternal-fetal interface, forming the placental barrier.^40^ Cell fusion of BeWo CTBs using forskolin is a widely adopted *in vitro* model for recapitulating trophoblast syncytialization.^41^; ^42^ Forskolin stimulates adenylyl cyclase, elevates intracellular cyclic AMP (cAMP), and activates the cAMP-PKA-CREB signaling axis, which transcriptionally drives cell fusion programs leading to syncytium formation. As the STB constitutes the maternal-facing layer of the placenta, establishing its susceptibility to MC-LR uptake is a key step in evaluating how this toxin may cross the maternal–fetal interface and affect placental/fetal health.

Despite emerging animal and *in vitro* studies demonstrating the effects of MC-LR on the placenta, the specific transport process that regulates MC accumulation and disposition in human placenta remains poorly defined. There remains a knowledge gap of direct evidence of MC accumulation and protein binding in human placental cells, which transporters mediate MC uptake, and how this transport is regulated by key transcriptional regulators HIF1A and CREB. Here, we present novel *in vitro* insights into the cellular entry of MC-LR into human immortalized trophoblasts. We evaluated MC-LR accumulation in widely used placental cell lines, including JAR and BeWo CTBs and HTR-8/SVneo extravillous trophoblasts (EVTs). We hypothesized that MC-LR undergoes active transport via OATP isoforms expressed in placental cells, and that uptake is influenced by physiological conditions such as oxygen concentration and cell fusion.

## Methods

### Chemicals

MC-LR (Cayman Chemicals Cat #10007188, Purity >95%) was purchased in large quantities to ensure the same lot number (Batch 0669496-1) was used across experiments. Cyclosporin A (cyc A, Cat #30024; Purity ≥95%) and fluorescein sodium salt (Cat #F6377, Purity 96%) were purchased from Sigma Aldrich (St. Louis, MO). All chemicals were reconstituted in 100% DMSO (D8418), with a maximum of 0.1% DMSO encountered by cells. All experimental treatment concentrations were diluted in Hanks Balanced Salt Solution (HBSS) (Gibco Cat #14170112). All reagents were carefully selected to prevent unintentional MC contamination.

### Cell Culture Maintenance

Human JAR (Cat #HTB-144), BeWo (Cat #CCL-98), and HTR-8/SVneo (Cat #CRL-3271) placental cell lines were purchased from ATCC (Manassas, VA). JAR and HTR-8/SVneo cells were cultured in phenol red-free RPMI1640 media (Gibco, Cat #11835-030) supplemented with 10% FBS (R&D Systems, Cat #S11550) and 1% PenStrep (Gibco, Cat# 15140122). BeWo cells were cultured in phenol red free DMEM/F:12 media (Gibco, Cat #11039-021) supplemented with 10% FBS and 1% PenStrep. JAR and BeWo choriocarcinoma cells are representative models for human placenta CTB cells, while HTR-8/SVneo cells are immortalized EVT cells.^43; 44^ All cells were maintained at 37°C in 5% CO_2_. Cell lines were maintained up to 20 passages for all experiments, and confirmed to be mycoplasma free (ATCC, Cat #30-1012K).

### RNA isolation and Real Time-quantitative PCR (RT-qPCR)

Following treatment, total RNA was extracted from cells using RNAzol (Sigma-Aldrich, Cat #R4533). Complementary DNA was synthesized from the isolated RNA with the High-Capacity cDNA Reverse Transcription Kit (Applied Biosystems by Thermo Fisher Scientific, Cat #4374967). Gene expression was quantified by RT-qPCR using gene-specific forward and reverse primers (Integrated DNA Technologies, Coralville, IA, USA, Supplemental Table 1) and PowerSYBR Green PCR Master Mix (Life Technologies, Cat #4367659). Amplification was performed on a ViiA7 Real-Time PCR System (Life Technologies). Threshold cycle (CT) values were analyzed using the ΔΔCT method, with beta-2-microglobulin (B2M) serving as the internal reference gene for normalization.

### Western Blot

Following treatment, cells were lysed with a 1X cell lysis buffer (Cat #89900, ThermoFisher) containing 25 mM Tris-HCl, pH 7.6, 150 mM NaCl, 1% NP40, 1% sodium deoxycholate, and 0.1% sodium dodecyl sulfate, spiked with 1% protease inhibitor cocktail (P8340, Sigma Aldrich, Burlington, MA). Pierce BCA assay (Thermo Scientific, Cat #23225) was run on all protein samples to determine total protein concentrations and adjusted to 15 μg protein for each sample. Protein was run on NuPAGE™ Bis-Tris Midi Protein Gels, 4 to 12%, 1.0 mm (Invitrogen, Cat #WG1402BOX). Protein was transferred onto a PVDF membrane (Thermo Scientific, Cat #88518) and transfer of proteins was confirmed with Ponceau S staining (Thermo Scientific, Cat #A40000279). All PVDF membranes were blocked with 5% fat-free dry milk in PBST for 2 hours. Primary antibodies were used to detect MC bound proteins MC (Cat #NBP2-89027, 1:200, Novus Biologicals, Littleton, CO), organic anion transporting polypeptide OATP1A2 (Cat #ab110392, 1:2000, Abcam, Cambridge, United Kingdom), OATP1B1 (Cat #ab312838,1:2000, Abcam), OATP1B3 (Cat #ab242368, 1:2000, Abcam), 2A1 (Cat #PA5-53083, 1:2000, Invitrogen), 3A1 (Cat #PA5-42457, 1:2000, Invitrogen), 4A1 (Cat #PA5-56535, 1:2000, Invitrogen), 5A1 (Cat #PA5-69280,1:2000, Invitrogen), PP2A (Cat #MA5-18060, 1:1000, Invitrogen Waltham, MA), β-Actin (Cat #ab8227, 1:500, Abcam), and GAPDH (Cat #ab9485 1:500, Abcam) and incubated overnight at HRP-conjugated secondary antibodies (anti -mouse or -rabbit, 1:2000, Sigma-Aldrich) were incubated for 1.5 hours. All primary and secondary antibodies were diluted in 2% milk in PBST. Chemiluminescent protein-antibody complexes were visualized using SuperSignal™ West Dura Extended Duration Substrate (Cat #34075, Thermo Fisher Scientific) and blot image captured on a Fluorchem Imager (ProteinSimple, Santa Clara, CA). Semi-quantitative analysis of relative protein expression was performed using Fiji package.^45^ Protein expression was normalized to β-Actin or GAPDH.

### MC-LR Uptake Experiments

JAR, BeWo, or HTR-8/SVneo cells were seeded at a concentration of 1.0 x 10^6^ cells/ml in Poly-D Lysine-coated 6-well plates (Gibco, Cat #A3890401). Cells were allowed to adhere overnight and treated at 80% confluency. Cells were treated with vehicle (0.1% DMSO), 0.1, 1, and 10 μM MC-LR in HBSS for 3 hours. Treatment was then aspirated, cells washed three times with ice cold PBS, cells lysed, and protein isolated for western blotting.

### Temperature-Dependent and OATP-Mediated Uptake of Fluorescent Substrate

JAR cells were suspended in 1.7ml microcentrifuge tubes at a concentration of 1.0×10^5^ cells/ml. JAR cells were pretreated with cyc A (vehicle or 1 μM) for 1 hour and then treated with fluorescein (vehicle or 1 μM) in HBSS for 4 time points between 10 and 40 minutes. Treatment was then aspirated, and cells were washed three times with ice cold PBS. Cells were then lysed, and lysate was then measured for fluorescence (Ex/Em: 494/515 nm) on microplate reader. A standard curve for fluorescein was used to calculate the concentration of fluorescein in cell lysates.

### Temperature-Dependent Uptake of MC-LR

JAR cells were seeded at a concentration of 1.0 x 10^6^ cells/ml in Poly-D Lysine-coated 6-well plates (Gibco, Cat #A3890401). Cells were allowed to adhere overnight and treated at 80% confluency. Cells were preincubated at either 4 or 37°C for 1 hour before treatment. Cells were then treated with MC-LR (vehicle or 1 μM) in HBSS and incubated at either 4 or 37°C for 1 hour. Treatment was then aspirated, cells washed three times with ice cold PBS, cells lysed, and protein isolated for western blotting.

### Inhibition of OATP Mediated Transport of MC-LR

JAR cells were seeded at a concentration of 1.0 x 10^6^ cells/ml in Poly-D-Lysine-coated 6-well plates (Gibco, Cat #A3890401). Cells were allowed to adhere overnight and treated at 80% confluency. JAR cells were treated with either vehicle (0.1% DMSO), cyc A (10 μM), MC-LR (1 μM), or the combination of cyc A (10 μM) + MC-LR (1 μM) for 1 hour incubated at either 4 or 37°C. Cells were pre-treated with cyc A (10 μM) for 1 hour before co-treatment with MC-LR (1 μM) to achieve potent inhibition of OATPs.^46^ Treatment was then aspirated, cells washed three times with ice cold PBS, cells lysed, and protein isolated for western blotting.

### Impact of Oxygen Tension on MC-LR Uptake

Hypoxia Cell Culture chambers (StemCell Technologies, Cat #27310) were used to maintain cell cultures at 3% and 8% O_2_ supplemented with 5% CO_2_ (Airgas, Radnor, PA). Ambient oxygen concentration (20% O_2_) in the cell culture incubator was used as the control. JAR cells were seeded at a concentration of 1.0 x 10^6^ cells/ml in Poly-D-Lysine-coated 6-well plates at either 3, 8, or 20% O_2_ for 24 hours. Cells were then treated with MC-LR (vehicle or 1 μM) for 1 hour. Treatment was then aspirated, cells washed three times with ice-cold PBS, cells lysed, and protein isolated for western blotting. Up-regulation of GLUT1, which is downstream of HIF1A, was measured to confirm hypoxic responses.^47^ OATP isoforms and PP expression were probed to detect changes in mRNA and protein expression at different oxygen tensions.

### Impact of Cell Fusion on MC-LR Uptake

BeWo cells were seeded at a density of 1.0 x 10^6^ cells/ml in Poly-D-Lysine-coated 6-well plates and allowed to adhere overnight. Cells were then treated with forskolin (vehicle or 20 μM) for 48 hours, after which media was collected to measure hCG production by ELISA (R&D systems, Cat #DY9034-05) as a marker for syncytialization.^41^ Cells were washed 3 times with PBS and treated with MC-LR (vehicle or 1 μM) for 3 hours. Treatment was then aspirated, cells washed three times with ice-cold PBS, cells lysed, and protein isolated for western blotting. Intracellular cAMP levels in cell lysate were measured by ELISA (Cayman Chemical, Cat #501040) to confirm activation of cell fusion pathways. OATP isoforms and PP expression were probed to detect changes in mRNA and protein expression after forskolin-stimulated cell fusion.

### Data Analysis

Data were presented as mean ± SD and analyzed using GraphPad Prism 9 (GraphPad Software Inc., La Jolla, CA). Outliers were identified and excluded by Grubb’s test. Data were normally distributed, and Student’s t-tests, one-way ANOVA with Dunnett’s post-hoc, or two-way ANOVA with Šídák’s multiple comparison post-hoc tests were performed to assess statistical significance (p < 0.05).

## Results

### Concentration-Dependent Uptake of MC-LR in Human Placental Cells

After *in vitro* MC-LR exposure of human placental cells for 6 hours, the MC-LR bound protein was observed at all MC-LR concentrations (0.1, 1, and 10 μM) with a concentration-dependent increase in JAR cells, while it was only detected at the highest concentration of 10 μM in BeWo cells. In HTR-8/SVneo cells, MC-LR bound protein was observed at concentrations of 1 μM or greater. These results suggest that there was a concentration-dependent increase in MC-LR accumulation that differed across human placental cell lines with more extensive accumulation in JAR and HTR-8/SVneo cells at lower concentrtaions (Figure 1).

**Figure 1.**
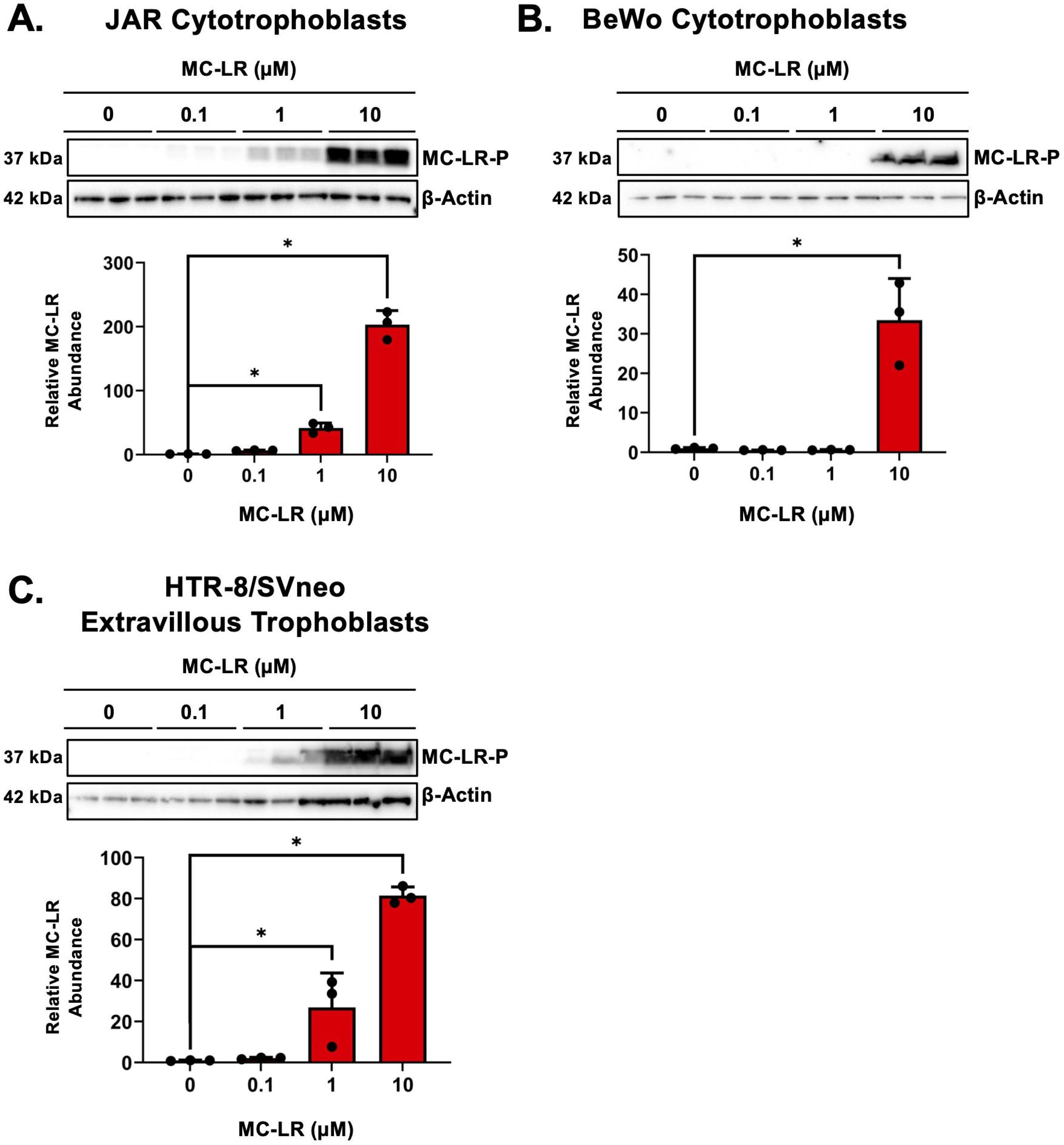
Concentration-Dependent Uptake of Microcystin-LR in Human Placenta Cell Lines. Three human placenta cell lines (cytotrophoblasts: JAR, BeWo, and extravillous trophoblasts: HTR-8/SVneo) were incubated with microcystin-LR (MC-LR) for 3 hours and then lysed before separation of proteins by SDS-PAGE and western blotting. MC-LR-bound proteins were visualized using an antibody that recognizes the 5 arginine microcystin congeners (LR, RR, YR, WR, FR). MW: MC-LR bound protein (MC-LR-P) corresponding to PP1/PP2A MW (37kDa), β-Actin (42kDa). Data represented as mean ± SD (n=3). One-way ANOVA with Dunnett’s multiple comparison posthoc, *p<0.05 compared to control (0 µM).

As variation in MC-LR uptake between placental cell lines may be explained by the different expression profiles of OATPs (Figure 2), multiple OATP isoforms (OATP1A2, 2A1, 2B1, 3A1, 4A1, 5A1) were next examined at both mRNA and protein levels in cells without MC-LR treatment. Lower threshold cycle (CT) values (*green)* indicate higher mRNA levels. Across the three placenta cells lines, there was enrichment of mRNA and protein expression for OATP1A2, OATP2B1, and OATP4A1. Little expression of OATP2A1, 3A1, or 5A1 was observed in the 3 cell lines with the exception of OATP3A1 mRNA in HTR-8/SVneo cells. Notably, mRNA and protein expression of OATP1B1 and 1B3 were not detected in JAR, BeWo, or HTR-8/SVneo cells (data not shown).

**Figure 2.**
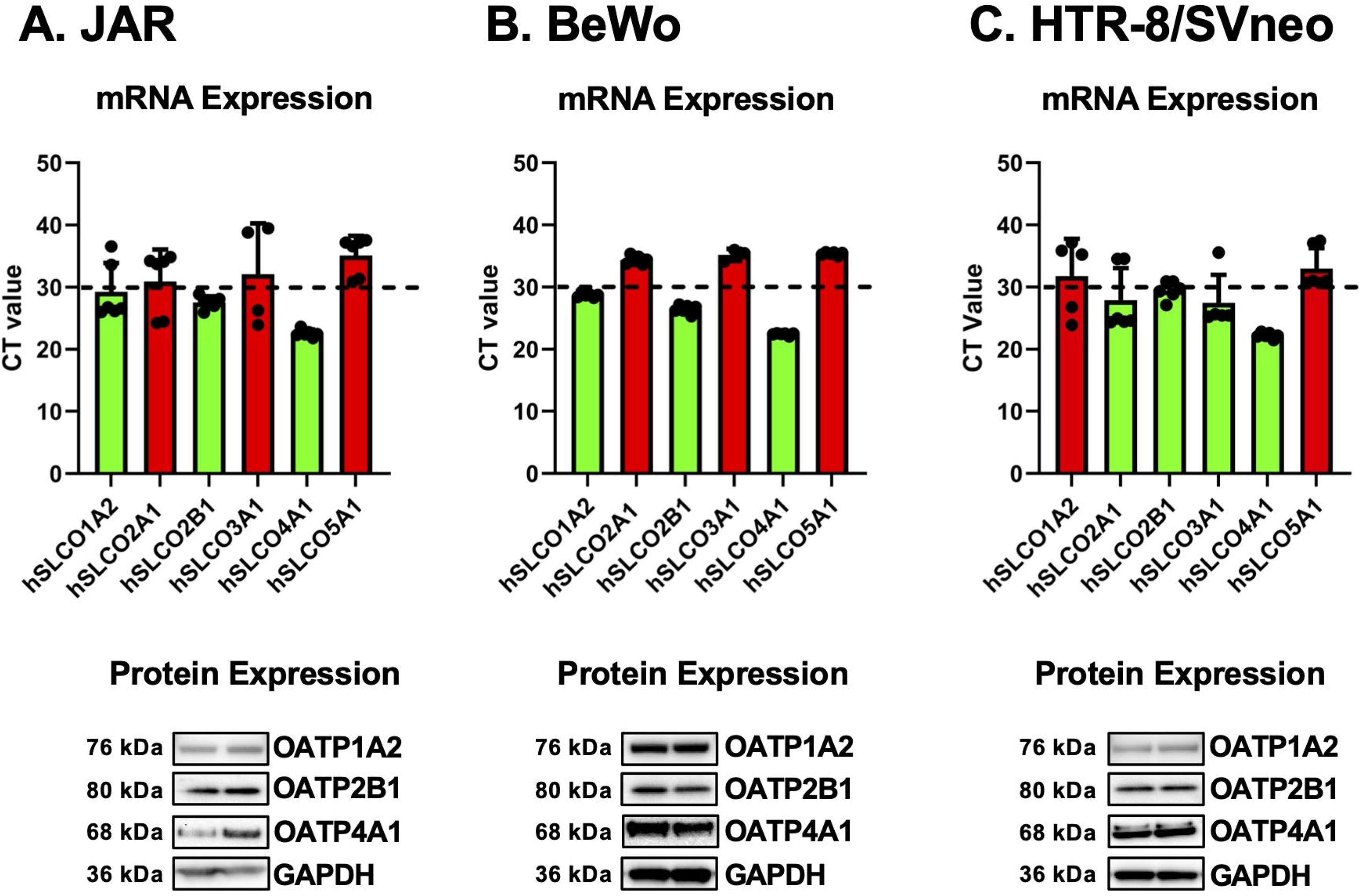
mRNA and Protein Expression of OATP Isoforms in Human Placenta Cell Lines. mRNA expression of hSLCO isoforms were measured by RT-qPCR (n=4) and protein (n=2) expression of OATP isoforms in (A) JAR, (B) BeWo, and (C) HTR-8/SVneo cells. MW: OATP1A2 (76 kDa), OATP2B1 (77 kDa) and OATP4A1 (68 kDa). mRNA expression was represented as raw Ct values, and a Ct threshold of 30 was used to identify lowly expressed genes (*red)* and enriched genes (*green*) in the three cell lines. Protein expression was normalized to GAPDH (36 kDa).

### Active Transport of MC-LR by OATP isoforms in Human Placental Cells

To define the mode of transport for MC-LR into JAR cells, we performed temperature-dependent uptake experiments and pharmacological inhibition studies targeting OATPs. At 37°C, both active and passive transport occur, while at 4°C passive transport predominates.^21; 48; 49^ There was time-dependent uptake of fluorescein at 37C with steady state reached by 20 min. JAR cells incubated at 4°C exhibited significantly reduced fluorescein uptake relative to 37°C, demonstrating temperature-dependent functional active transport in JAR cells (Figure 3A). In addition, JAR cells were pretreated with the general OATP inhibitor cyc A (10 μM) for 1 hour and then co-treated with the OATP substrate, fluorescein (1 μM). The uptake of fluorescein was inhibited 45% by Cyc A, confirming OATP activity in JAR cells (Figure 3A).

**Figure 3.**
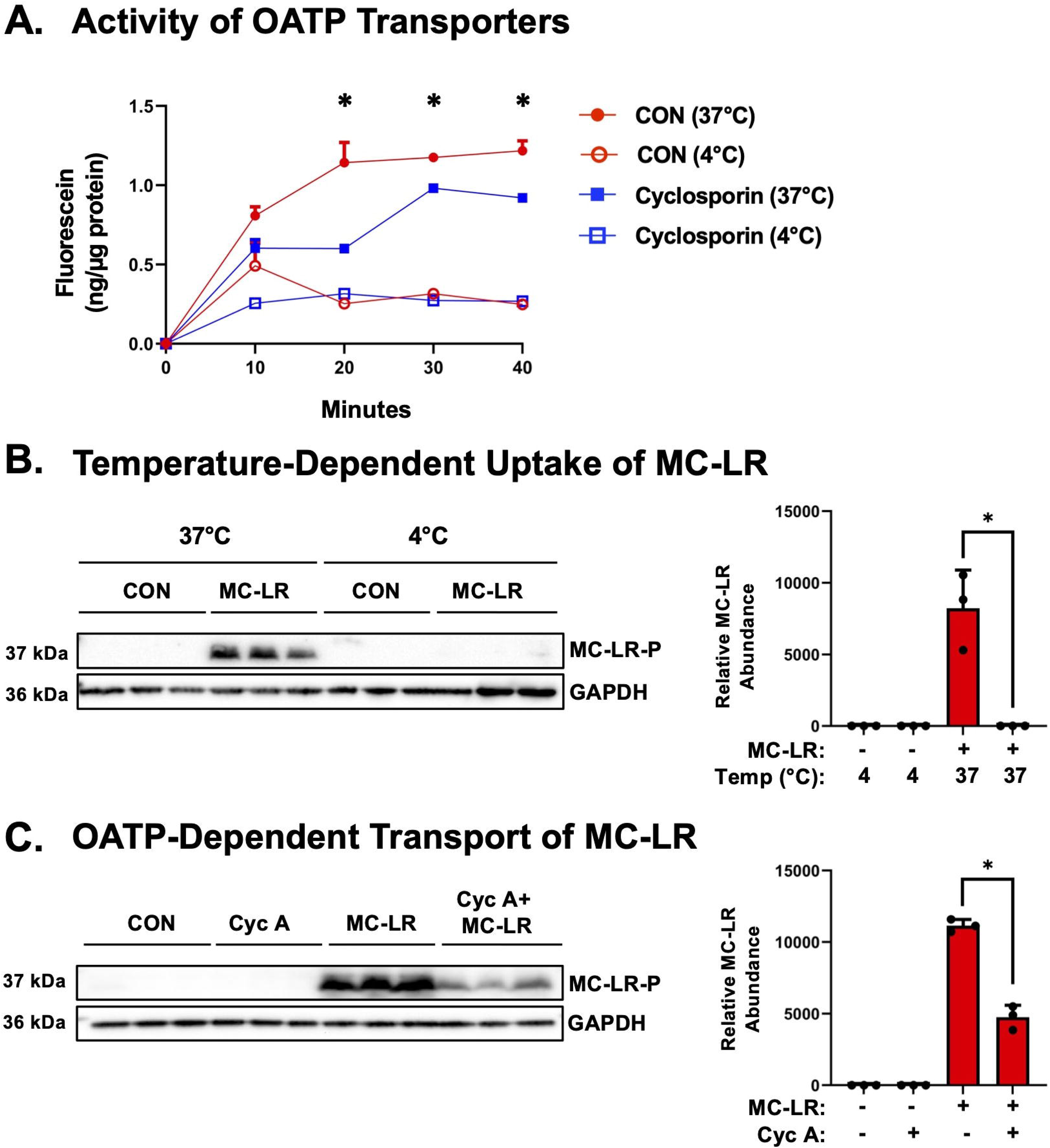
OATP-Dependent Uptake of Fluorescein and Microcystin-LR in Human Placenta Cells. (A) Fluorescein is a fluorescent substrate for multiple OATP isoforms. Active and passive transport occur at 37°C, with only passive transport occurring at 4°C. JAR cytotrophoblast cells were pretreated with 10 μM cyc A, a general OATP inhibitor, for 1 hour and then treated with 1 μM fluorescein for up to 30 min. Fluorescence was measured at 494 nm ex/515 nm em and compared to a standard curve. *p<0.05 compared to cyc A-treated cells at 37°C. (B) JAR cells were incubated at either 37°C (active and passive transport) or 4°C (passive transport only) with 1 μM MC-LR for 3 hourrs. *p<0.05 comparing 1 μM MC-LR uptake at 37°C. (C) JAR cells were pre-treated with 10 μM cyc A, a general OATP inhibitor, for 1 hour and then treated with 1 μM MC-LR for 40 min. Cells were lysed and probed for MC-LR-bound proteins. Uptake of MC-LR and subsequent binding was reduced by inhibiting OATP transporters. Data represented as mean ± SD (n=3-4). Two-way ANOVA with Dunnett’s multiple comparison posthoc, *p<0.05 comparing MC-LR vs Cyc A + MC-LR.

The next set of experiments were performed with MC-LR. At 37°C, there was significantly increased uptake of MC-LR in JAR cells compared with the vehicle control group, while JAR cells incubated at 4°C did not exhibit uptake of MC-LR (Figure 3B). In addition, there was a 57% reduction in MC-LR (1 μM) uptake in cyc A-treated cells compared to controls (Figure 3C). Collectively, these results demonstrate the active transport of MC-LR by OATP-mediated transport in human placental cells.

### OATP-Mediated Transport and MC-LR Uptake in Human Placenta Cells at Low Oxygen Concentrations

JAR cells were incubated at 20, 8, or 3% O_2_ for 48 hours, and the mRNA expression of genes associated with exposure to low oxygen concentrations were evaluated. Compared to 20% O_2_, cells exposed to 8% or 3% O_2_ had increased mRNA expression of GLUT1, VEGFR1, and HO-1 (only at 3%), but no change in HIF1A mRNA was observed (Figure 4A). Next, OATP transport activity was assessed in JAR cells exposed to 20%, 8%, and 3% O_2_. Compared to cells grown at 20% O_2,_ the uptake of fluorescein was enhanced by 80% at 8% O_2_ and 111% at 3% O_2_ (Figure 4B). Based on these results, we hypothesized that increased OATP expression during hypoxia would lead to increased uptake of MC-LR. Unexpectedly, compared to 20% O_2_, MC-LR bound protein was decreased by 52% and 72% in JAR cells exposed to 3% and 8% O_2,_ respectively (Figure 4C). To further investigate the underlying mechanism, the mRNA and protein expression of OATPs and PPs were examined (Figure 4D and Supplemental Figure 1). The results of RT-qPCR revealed a slight up-regulation of OATP2B1 mRNA expression with hypoxia, although the difference was not statistically significant. However, western blotting revealed that only the expression of OATP4A1 was significantly up-regulated 6.9-fold with lower oxygen concentrations of 3% and 8% O_2_, while OATP2B1 tended to be increased by 2.8-fold (p-value= 0.14) and OATP1A2 protein was decreased by 60% at 3% O_2_. Somewhat surprisingly, protein phosphatase PP2A mRNA and protein expression were significantly reduced by up to 90% at 8% and 3% O_2_.

**Figure 4.**
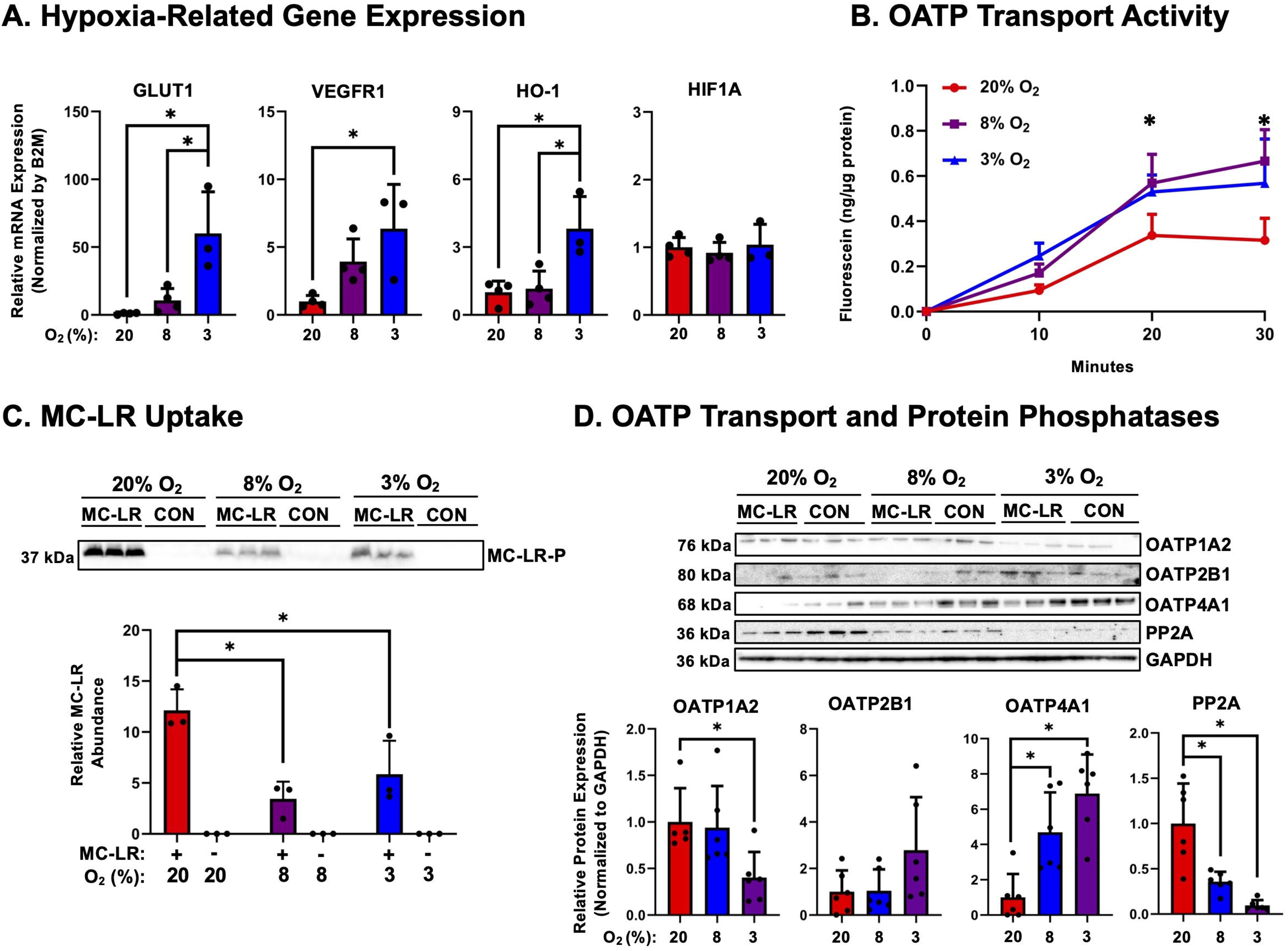
Transport of Microcystin-LR into Human Placenta Cells at Different Oxygen Concentrations. JAR cells were exposed to oxygen concentrations of 20%, 8%, and 3% O_2_ for 24 hours. (A) Expression of GLUT1, VEGFR1, HO-1, and HIF1A mRNAs were quantified as positive control responses to low O_2_ concentrations. *p<0.05 comparing to 20% O_2_ control. (B) JAR cells were treated with fluorescein (0 or 1 μM) for time points between 0-30 minutes. Cells were then lysed and lysate measured for fluorescence (ex/em:494/515). (C) JAR cells were treated with MC-LR (0 or 1 μM) for 1 hour, and MC-LR bound protein was quantified by western blotting. (D) Protein expression of OATP isoforms (1A2, 2B1, and 4A1) and protein phosphatases (PP2A) in JAR cells at oxygen concentrations of 20, 8, and 3% O_2_ were measured. Data represented as mean ± SD (n=3-6). Two-way ANOVA with Šídák’s multiple comparisons test. *p<0.05 compared to JAR cells at 20% O_2_.

### OATP-Mediated Transport and MC-LR Uptake in Human Placenta Cells After Forskolin Stimulation of Cell Fusion

We next treated BeWo cells with forskolin, an established approach to stimulate intracellular cAMP levels and induce trophoblast cell fusion. ELISA results showed that BeWo cells treated with forskolin (20 μM) had a significant increase in intracellular cAMP over 200% (Figure 5A). Furthermore, there was a corresponding increase in hCG secretion (81%, Figure 5A) after forskolin treatment, a marker of increased placental function observed after cell fusion. Compared to vehicle-treated cells, BeWo cells treated with forskolin exhibited increased uptake of the OATP substrate fluorescein (1 μM) by up to 237% over 7.5 minutes (Figure 5B) and MC-LR by 263% (Figure 5C). The expression of OATPs or PPs was next examined to determine whether their expression influenced MC-LR uptake after forskolin treatment (Figure 5D and Supplemental Figure 2). After forskolin treatment, the mRNA expression of OATP1A2, a known transporter for MC-LR, was reduced with little to no change in protein expression. While there was increased expression of OATP2B1 mRNA in forskolin-treated cells, protein levels declined by 50%. Interestingly, while the mRNA expression of OATP4A1 did not change significantly, its protein expression increased by 70%. Additionally, there were moderate increases in the mRNA expression of PP1A and PP2A in forskolin-treated cells, however, there was no significant change at the protein level.

**Figure 5.**
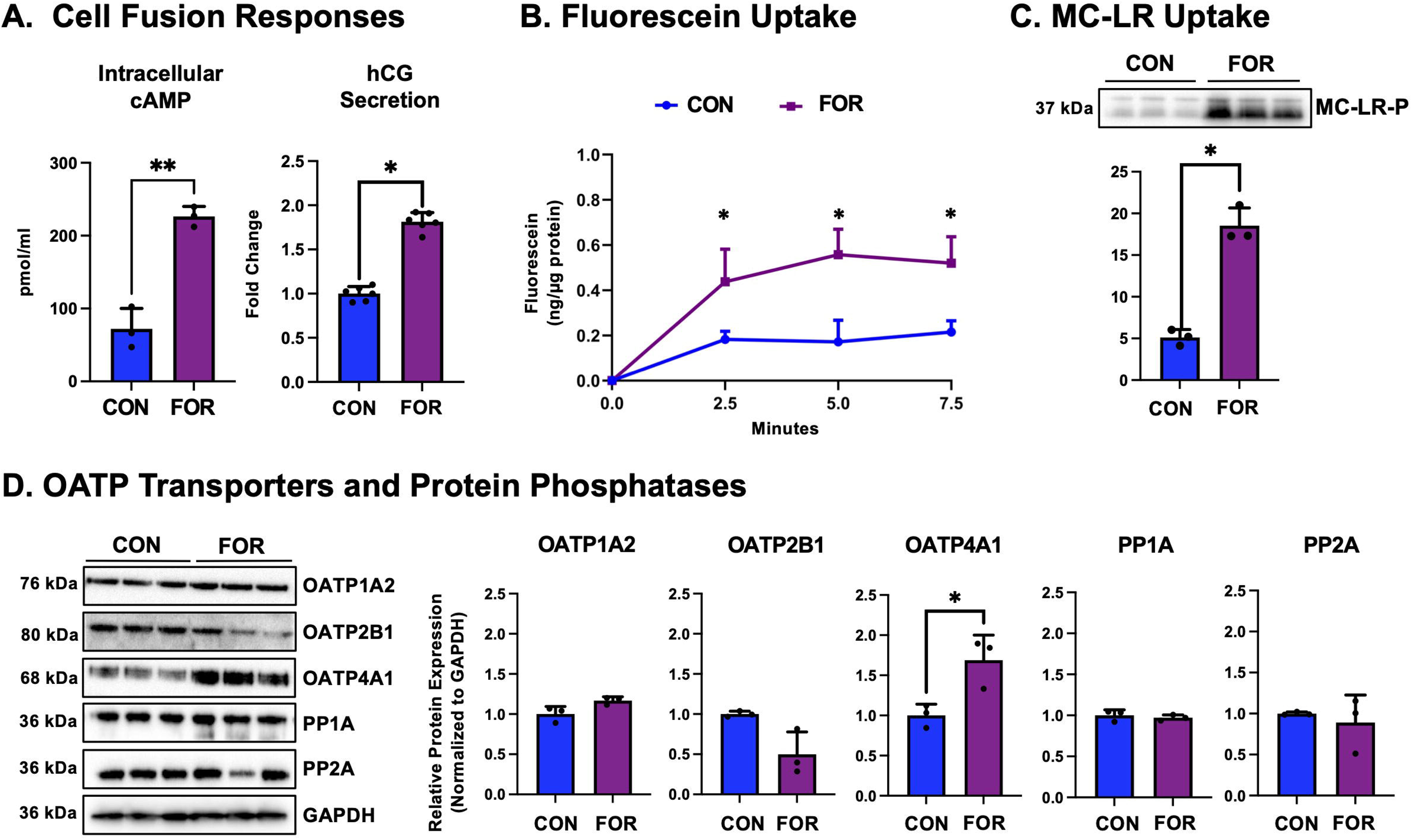
Transport of Microcystin-LR into Human Placenta Cells Stimulated to Undergo Fusion. BeWo cells were treated with either 0 or 20 μM forskolin (FOR) for 48 hours to stimulate fusion. (A) Intracellular cAMP and HCG secretion were quantified by ELISA. (B) Fluorescein uptake (0 or 1 μM) into control (CON) and forskolin (FOR)-treated BeWo cells was measured over 10 minutes (ex/em: 494/515 nm) and compared to a standard curve. (C) MC-LR uptake (0 or 1 μM) into control (CON) and forskolin (FOR)-treated BeWo was quantified after 1 hour. (D) Protein expression of OATP isoforms (1A2, 2B1, and 4A1) and protein phosphatases (PP1A and PP2A) were measured in control (CON) and forskolin (FOR)-treated BeWo cells. Data represented as mean ± SD (n=3-6) Statistical significance was assessed using an unpaired t-test, *p<0.05 compared to control.

## Discussion

Our research posits that MCIZLR uptake in placental cells occurs through a temperatureIZsensitive, transporterIZmediated process, most likely involving OATP isoforms uniquely expressed in the placenta. JAR cells were used as the model trophoblast cell line for MC-LR transport experiments, as they showed the most sensitive uptake of MC-LR. The OATP expression profiles of JAR, BeWo, and HTR-8/SVneo cells identify OATP1A2, OATP2B1, and OATP4A1 as the placental isoforms most likely to mediate MC-LR transport. OATP1A2 has previously been demonstrated to transport MC-LR, indicating that it may also contribute MC-LR uptake in immortalized trophoblast cell lines.^49^ Pan OATP inhibitors, including cyc A, have been shown to inhibit OATP transport.^46^ The significant reduction of MC-LR uptake in cyc A-treated JAR CTBs demonstrates that human placenta cells possess OATPs-mediated MC-LR transport.

Across the three human trophoblast cell lines, cytotoxicity was assessed by propidium iodide staining and lactate dehydrogenase release, which showed no significant toxicity after 24 hours of exposure to 0.1–10 μM MCIZLR (data not shown), aligning with findings in primary villous CTBs that reported no morphological toxicity (cell rounding or detachment) below 25 μM MCIZLR after 24 hour treatment.^30^ This resistance to high concentrations of MC-LR in CTBs contrasts with hepatocyte models, where MCIZLR can trigger apoptosis at low concentrations less than or equal to 2 μM MC-LR.^50^ Studies in HepG2 liver cells further suggest that cytotoxicity may be contextIZdependent, with enhanced sensitivity under hypoxic conditions, likely due to increased expression of OATPs and exacerbated oxidative stress.^51^ These findings support the idea that susceptibility to MCIZLR is strongly influenced by cellular redox state and transporter activity, which may explain the low cytotoxicity of trophoblasts observed under atmospheric conditions.

Incorporation of physiologically-relevant O_2_ levels observed during early and late trimesters of pregnancy *in vitro*, revealed significant changes in OATP transporter expression and protein phosphatases. At lower oxygen concentrations, which are observed early in pregnancies and during pathological conditions^52^, there was increased uptake of the OATP substrate fluorescein but decreased uptake of MC-LR. We next examined OATP and PP expression at both mRNA and protein levels to clarify the molecular basis of this effect. OATP1A2, a known transporter of MC-LR, exhibited reduced mRNA and protein expression at 3% O_2_, potentially reducing the amount of MC-LR uptake. There was modest increased expression of OATP2B1 mRNA and protein expression at 3% O_2_, however, OATP2B1 has previously been reported to not significantly contribute to the transport of MC-LR.^49^ Most strikingly, there was up-regulation in OATP4A1 protein expression, explaining the increased uptake of the OATP substrate fluorescein. The reduction in observed MC-LR bound protein most likely coincides with the reduced PP2A target protein expression at 8 and 3% O_2_, suggesting a larger unbound MC-LR fraction in trophoblasts under hypoxia. This aligns with other studies investigating MC-LR toxicity during hypoxia, finding increased oxidative stress at low oxygen concentrations.^51^ Overall, our findings indicate that low oxygen tension differentially regulates OATP and PP expression, with OATP4A1 likely mediating MC-LR transport in JAR CTBs. Previous research from our laboratory found that low O_2_ concentrations differentially regulate the expression of placental solute carriers across two experimental human placental models, including BeWo CTBs and term placenta villous explants.^53^ Notably, there was a significant decrease in *SLCO*/OATP4A1 mRNA expression in BeWo cells, but an observed trending increase in term placental villous explants incubated at low oxygen concentrations. There is well-established evidence supporting a HIF1A/OATP regulatory axis, whereby hypoxia activates HIF1A, leading to transcriptional upregulation of downstream OATP transporters and consequently enhancing substrate uptake in cancer cells.^54–56^ While we likely missed the window of OATP4A1 mRNA up-regulation as levels were only measured at 24 hours, these results indicate that the regulation of uptake transporters, such as OATPs, is influenced by low oxygen concentrations.

Finally, because STBs are the placental cell type directly in contact with the maternal circulation, establishing whether MC-LR can enter these cells is of critical importance. Using forskolin-treated BeWo cells as an *in vitro* model of STBs, we observed a marked increase in intracellular cAMP following 48 hours of treatment, an indicator of activation of signaling pathways that regulate trophoblast cell fusion. Moreover, there was a 80% increase in hCG secretion, a hallmark of increased placenta function associated with trophoblast fusion (Figure 5A). Functional uptake assays revealed that forskolin stimulation significantly enhanced the accumulation of both fluorescein and MC-LR, with MC-LR uptake elevated by 263% compared to control BeWo cells (Figure 5B–C). These findings indicate that MC-LR entry is enhanced in syncytialized cells. To explore the molecular mechanism, we quantified OATP and PP mRNA and protein expression (Figure 5D–E). OATP4A1 exhibited no change in mRNA levels yet demonstrated the most pronounced up-regulation at the protein level after forskolin stimulation. Taken together, these results suggest that regulation of OATPs, particularly OATP4A1, is influenced by syncytialization, leading to enhanced uptake of MC-LR in STBs, highlighting a potential mechanism by which maternal exposure could directly impact the syncytial barrier. Previous research has found that forskolin stimulated cell fusion of BeWo CTBs leads to up-regulation of the SLC transporter, OAT4, expression and function, while OATP2B1 expression was unchanged or undetectable.^57^ However, treatment of HRP-1 rat trophoblast cells with 100 μM forskolin or 250 μM 8-Br-cAMP for 3 hours significantly increased uptake of the OATP substrate [³H]estrone sulfate, and this uptake was further enhanced by 3-isobutyl-1-methylxanthine, which sustains intracellular cAMP levels by inhibiting cyclic nucleotide phosphodiesterase.^58^ These data suggest that both increased cell fusion and elevated intracellular cAMP levels may impact membrane transporter expression and function, potentially influencing MC-LR transport in STBs.

The glutamine–glutamate concentration gradient across the STB provides a strong driving force for OATP-mediated transport, as high intracellular glutamate levels (5–10 mM) enable exchange of organic anions across both the apical and basal membranes.^59^ This glutamine–glutamate cycling not only supports uptake of essential substrates such as estrone sulfate, thyroid hormones, and dehydroepiandrosterone sulfate, but also creates a pathway by which xenobiotics may enter the placenta. If coupled to the intracellular glutamate gradient, OATP4A1 (apical and basolateral) and OATP2B1 (basolateral) can drive secondary active uptake of organic anions, meaning they could facilitate maternal-to-placental and placental-to-fetal transfer of xenobiotics and toxins such as MC-LR. Whereas OATP2A1 (apical) does not appear glutamate-coupled and function more narrowly on the clearance of prostaglandins, we speculate it is less likely to be involved in MC-LR accumulation.^60^ In this way, glutamate coupling could facilitate OATP4A1 and OATP2B1 to act as entry routes for MC-LR into the STB and potentially the fetus. In the context of pathological pregnancy, reduced glutamate transporter activity has been observed in fetal growth restriction, suggesting that disruption of this gradient may impair placental transporter function.^61^ Beyond its role in transport, glutamate is central to antioxidant defense as the γIZglutamyl backbone of glutathione (GSH), which is synthesized with cysteine and glycine.^62^; ^63^ GSH efflux is tightly coupled to organic anion exchange and driven by the steep intracellular-to-extracellular GSH gradient. Studies using Oatp1 or Oatp2IZexpressing *Xenopus laevis* oocytes demonstrated that OATPs mediate bidirectional organic solute transport in a GSHIZsensitive manner, where high intracellular GSH or its conjugates stimulate taurocholate and digoxin uptake, while OATPs also facilitate GSH efflux.^64; 65^ Similar interactions have been observed in HepG2 cells, where depletion of intracellular GSH suppressed taurocholate uptake, whereas the presence of extracellular taurocholate and other bile acids enhanced GSH efflux.^66^ Together, these findings highlight that the intracellular GSH pool and its electrochemical gradient not only energize organic anion transport but also suggest that GSH conjugation status may influence the disposition of MC-LR by modulating its transport efficiency and intracellular accumulation. Omitting this interaction risks overlooking critical determinants of MC-LR bioavailability and toxicity, underscoring the need for future studies to examine how glutathione levels and conjugation status affect OATP-mediated uptake across isoforms.

Our current study has notable limitations including the fact that the cell lines used in these studies possess multiple OATP isoforms.^67^ Transport studies with overexpressing cells are needed to compare the kinetics of MC-LR transport across OATP isoforms. In addition, cytotoxicity studies will be performed in primary CTBs, after spontaneous syncytialization, to elucidate MC-LR uptake into native tissue and potential consequences to placental health and function.

## Conclusion

Our study demonstrates that MC-LR enters human trophoblasts by active OATP transporters in a concentration-dependent manner. This uptake is most likely mediated through OATP4A1, which is highly expressed in immortalized human trophoblast cell lines. During conditions of hypoxia, such as during early or pathological pregnancy, there is a significant up-regulation of OATP transporters, leading to higher uptake of MC-LR. However, due to reduced PP2A target protein expression, there is a decreased MC-LR bound protein fraction (PP inhibition), and likely a higher free unbound MC-LR fraction available in the cells. Moreover, there was significant enhanced uptake of MC-LR in fused CTBs, a model of the placental syncytial barrier. These insights address that accumulation and disposition of MC-LR in human placenta is influenced by low oxygen tension and trophoblast cell fusion.

## Acknowledgments

This work was supported by the National Institute of Environmental Health Sciences [Grants T32ES007148, R01ES029275, P30ES005022, R01ES032144, R25ES020721], National Institute of Child Health and Human Development [Grant UC2HD113039 and UC2HD113039-02S1] and National Center for Advancing Translational Sciences [Grant TL1TR003019], the Amnion Foundation, and the Dr. Gary and Mrs. Janis Grover Fellowship.

## Abbreviations

(B2M): beta-2-microglobulin
(cAMP): cyclic AMP
(cyc A): cyclosporin A
(CTBs): cytotrophoblasts
(EVTs): extravillous trophoblasts
(GLUT): glucose transporter
(GSH): glutathione
(HABs): harmful algal blooms
(HBSS): Hanks Balanced Salt Solution
(HIF): hypoxia-inducible factor
(MC-LR): microcystin-LR
(OATP): organic anion transporting polypeptide
(PP1/2A): protein phosphatase 1/2A
(SLCO): Solute Carrier superfamily
(STB): syncytiotrophoblast
(CT): threshold cycle

