## Supplementary figures and images for "Transporter-Mediated Uptake of Microcystin-LR in Human Trophoblasts: Regulation By Oxygen Concentration and Cell Fusion"

### Supplemental Figure 1

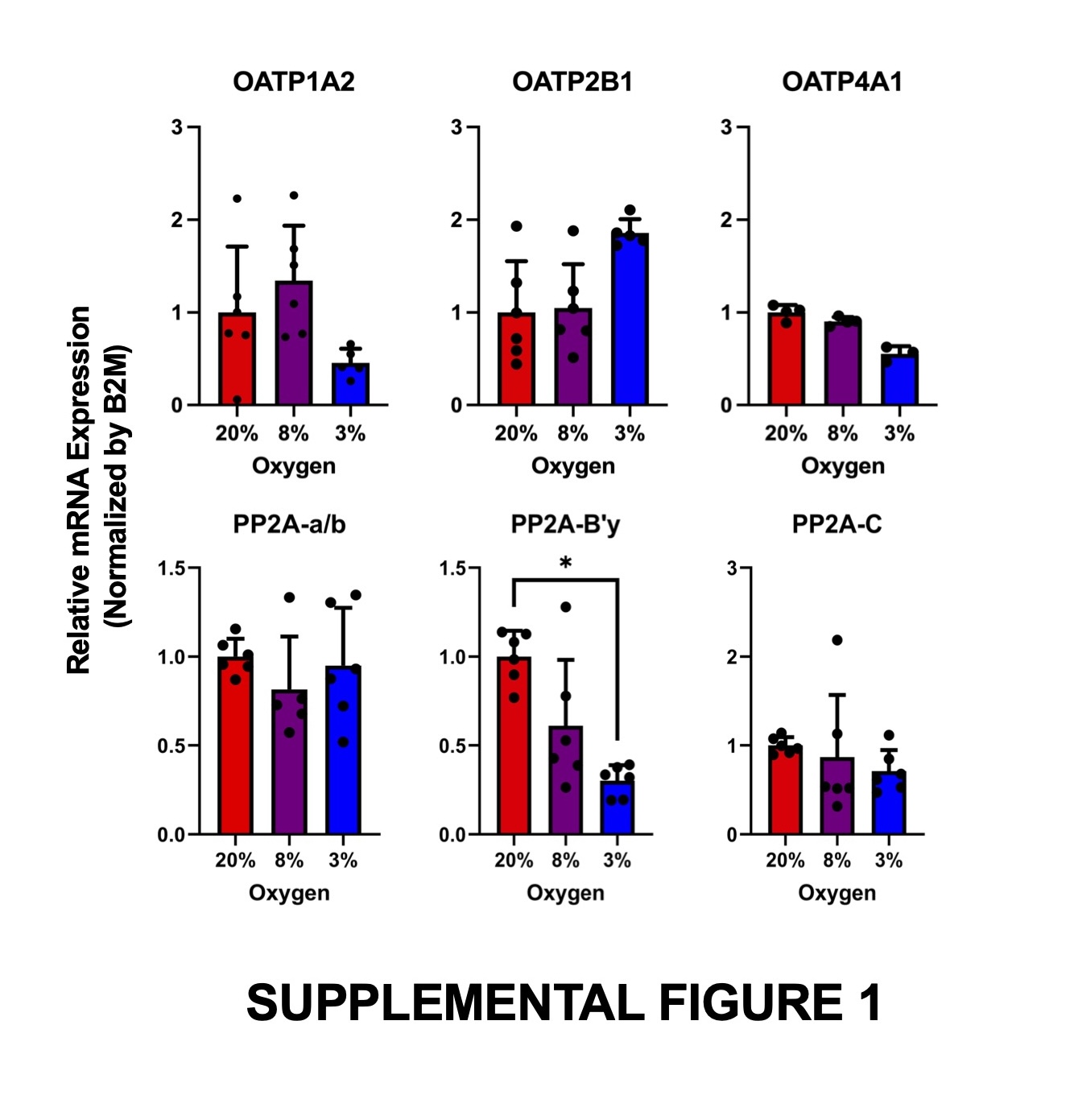

### Supplemental Figure 2

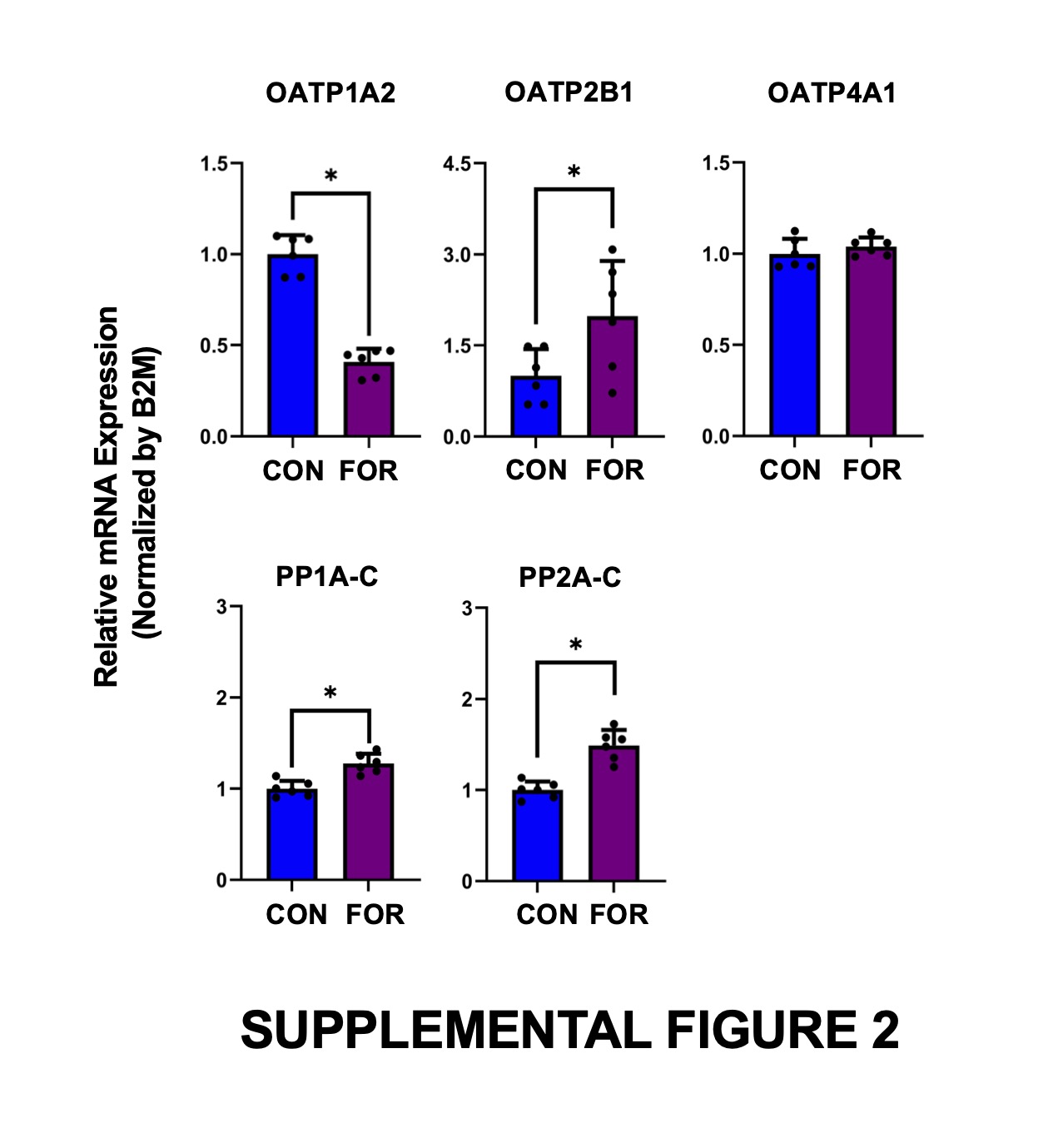
