## Supplemental Materials for "Transporter-Mediated Uptake of Microcystin-LR in Human Trophoblasts: Regulation By Oxygen Concentration and Cell Fusion"

**Affiliations:**

**Supplemental Figure 1. mRNA expression of OATPs and PPs in Human Placenta Cells at Different Oxygen Concentrations.** JAR cells were exposed to oxygen concentrations of 20%, 8%, and 3% O_2_ for 24 hours. mRNA expression of hSLCO/OATP isoforms (1A2, 2B1, and 4A1) and protein phosphatases (PP2A: -a/b (scaffold subunit), -B’y (regulatory subunit), and -C (catalytic subunit)) in JAR cells at oxygen concentrations of 20, 8, and 3% O_2_ were measured_._ Data represented as mean ± SD (n=6). Two-way ANOVA with Šídák's multiple comparisons test. *p<0.05 compared to JAR cells at 20% O_2_.

**Supplemental Figure 2. mRNA expression of OATPs and PPs in Human Placenta Cells Stimulated to Undergo Fusion.** BeWo cells were treated with either 0 or 20 μM forskolin (FOR) for 48 hours to stimulate fusion. mRNA expression of hSLCO/OATP isoforms (1A2, 2B1, and 4A1) and protein phosphatases (PP1A-C (catalytic subunit) and PP2A-C (catalytic subunit)) in control (CON) and forskolin (FOR)-treated BeWo cells. Data represented as mean ± SD (n=6). Statistical significance was assessed using an unpaired t-test, *p<0.05 compared to control.

**Supplemental Table 1. Primer Sequences for RT-qPCR.**

| **Gene Name** | **Primer Name** | **Forward (5'-3’)** | **Reverse (3’-5’)** |
| --- | --- | --- | --- |
| SLCO1A2 | OATP1A2 | TCTTCCTGACAAGATGGTGCT | TCTTCAGGGTGTTCCAAGCTA |
| SLCO1B1 | OATP1B1 | TGAACACCGTTGGAATTGC | TCTCTATGAGATGTCACTGGAT |
| SLCO1B3 | OATP1B3 | CCGTATTTTTTGGAAGGGTCTAC | TTCTTTCATTGTCCGATGCC |
| SLCO2A1 | OATP2A1 | CTGATAGCCTGCATCTCCCA | AGGAGTGGTCAATGGTGAGG |
| SLCO2B1 | OATP2B1 | TGATTGGCTATGGGGCTATC | CATATCCTCAGGGCTGGTGT |
| SLCO3A1 | OATP3A1 | CCTGCGTCCTCTACGACAAT | GGGTCAGAGTAGAGGCAAAGA |
| SLCO4A1 | OATP4A1 | GCGGAAATGCACCAGTTGAAGG | GGTTCTTCAGCAGGAGCCAGAT |
| SLCO5A1 | OATP5A1 | GGAGAGACCTTTTGCACTGG | TCACGTTGTACTCCCAGCAA |
| PP1A | Protein phosphatase 1A | GCTGGAAGGTGACATACACG | GATCTTATAGGCCAGCAGCAG |
| PP2A-C | Protein phosphatase 2A (PP2A-C) - catalytic | TCGTTGTGGTAACCAAGCTG | AACATGTGGCTCGCCTCTAC |
| PP2A-α/β | Protein phosphatase 2A=Aα/β (regulatory) | GCTTCAATGTGGCCAAGTCT | GGTCCTGGGTCAGCTTCTCT |
| PP2A-B’y | Protein phosphatase 2A-B’y (scaffold) | ACAGTGAAGGACGAGGCTCA | CTTCCAAGGCTTTCTTGGTG |
| B2M | beta-2 microglobulin | TCGCTCCGTGGCCTTAGCTG | CAATGTCGGATGGATGAAACCCAG |
